# Genome-wide scan and fine-mapping of rare nonsynonymous associations implicates intracellular lipolysis genes in fat distribution and cardio-metabolic risk

**DOI:** 10.1101/372128

**Authors:** Luca A. Lotta, Liang Dong, Chen Li, Satish Patel, Isobel D. Stewart, Koini Lim, Felix R. Day, Eleanor Wheeler, Craig A. Glastonbury, Marcel Van de Streek, Stephen J. Sharp, Jian’an Luan, Nicholas Bowkera, Martina Schweiger, Laura B. L. Wittemans, Nicola D. Kerrison, Lina Cai, Debora M. E. Lucarelli, Inês Barroso, Mark I. McCarthy, Robert A. Scott, Rudolf Zechner, John R. B. Perry, Vladimir Saudek, Kerrin S. Small, Stephen O’Rahilly, Nicholas J. Wareham, David B. Savage, Claudia Langenberg

## Abstract

Difficulties in identifying causal variants and genes underlying genetic associations have limited the translational potential of genetic studies of body fat distribution, an important, partly-heritable risk factor for cardio-metabolic disease. Rare variant associations facilitate fine-mapping of causal alleles, but their contribution to fat distribution is understudied. We performed a genome-wide scan of rare nonsynonymous variants for body mass index-adjusted waist-to-hip-ratio (BMI-adjusted WHR; a widely-used measure of fat distribution) in 450,562 European ancestry individuals, followed by systematic Bayesian fine-mapping at six genome-wide (p<5×10^−08^; main-analysis) and two subthreshold signals (significant at a Bonferroni-corrected p<1.3×10^−06^). We found strong statistical evidence of causal association for nonsynonymous alleles in *CALCRL* (p.L87P, p_conditional_=5.9×10^−12^; posterior-probability of association [PPA]=52%), *PLIN1* (p.L90P, p_conditional_=5.5×10^−13^; PPA>99%), *PDE3B* (p.R783X, p_conditional_=6.2×10^−15^; PPA>99%), *ACVR1C* (p.I195T; p_conditional_=5.4×10^−12^; PPA>99%), and *FGF1* (p.G21E, p_conditional_=1.6×10^−07^; PPA=98%). Alleles at the four likely-causal main-analysis genes affected fat distribution primarily via larger hip-rather than smaller waist-circumference and six of nine conditionally-independent WHR-lowering index-variants were associated with protection from cardiovascular or metabolic disease. All four genes are expressed in adipose tissue and have been linked with the regulation of intracellular lipolysis, which controls fat retention in mature cells. Targeted follow-up analyses of key intracellular-lipolysis genes revealed associations for a variant in the initiator of intracellular lipolysis *PNPLA2* (p.N252K) with higher BMI-adjusted-WHR and higher cardio-metabolic risk. This study provides human genetic evidence of a link between intracellular lipolysis, fat-distribution and its cardio-metabolic complications in the general population.

## Introduction

Body fat distribution is a major risk factor for cardiovascular and metabolic disease independent of obesity (1–5), but the mechanistic foundations of this link are poorly understood. Several common variants associated with fat distribution (6) or with compartmental fat deposition (7, 8) are located in regulatory regions near genes that are highly expressed in adipose tissue. While consistent with a biologically plausible role of adipocyte function in fat distribution, the translational potential of these observations is limited by challenges in inferring causal genes and mechanisms underlying associations of non-coding genetic variants. In contrast, the study of low-frequency nonsynonymous alleles has catalyzed translation from gene identification to therapeutic drug development, as illustrated by associations in *PCSK9* (9, 10), *LPA* (11), *APOC3 (12, 13)* or *ANGPTL3 (14–16)* with lipid phenotypes leading to rapid drug development for cardiovascular prevention (15, 17–22).

Previous genome-wide association studies of fat distribution in ~225,000 people have focused mostly on common variants (6). Rare variants, defined by the 1000 Genomes Project by minor allele frequency (MAF) below 0.5% (23), are usually population-specific (23) and difficult to impute (24), and hence their study requires large, homogeneous samples and direct genotyping. Their contribution to fat distribution remains understudied. A critical advantage of studying rare variants is that they represent mutational events which occurred more recently in a population (23), so that they tend to occur on long haplotypes together with more common variation with which correlation is low (25). This facilitates statistical fine-mapping aimed at identifying causal variants and distinguishing scenarios where the rare variant is causal rather than just a “passenger” in the association signal (25). Causal nonsynonymous variants in a gene provide a strong link between gene and phenotype and also a “genetic model” for functional studies aimed at understanding the underlying mechanism of association.

To exploit these properties, we conducted a genome-wide discovery scan of the association of rare nonsynonymous variants with body mass index-adjusted waist-to-hip ratio (BMI-adjusted WHR; a widely-used measure of body fat distribution (5, 6)) in 450,562 European ancestry individuals. We then conducted systematic analyses of genomic context to distinguish likely-causal from non-causal associations. The aim was to identify variants, genes and pathways implicated in the regulation of fat distribution in the general population.

## Results

### Genome-wide scan of rare nonsynonymous variants and fine-mapping at identified loci

We conducted a genome-wide association scan of 37,435 directly-genotyped, rare (MAF <0.5%) nonsynonymous variants with BMI-adjusted WHR in 450,562 European ancestry participants of UK Biobank (**SI Appendix Notes S1-S2, Tables S1-S2 and Fig. S1**). There was evidence of modest inflation of signal (λ=1.045), consistent with polygenic contributions to BMI-adjusted WHR (**SI Appendix Fig. S2**). In the main analysis, we identified six associations at the genome-wide level of statistical significance (p<5×10^−08^) in genes at least 1 Mb apart of each other (**SI Appendix Table S3 and Fig. S2**). In addition to the main analysis, we also identified two additional signals in a women-specific analysis (p<5×10^−08^ in a women-only secondary analysis; **SI Appendix Table S3**) and two additional signals that met a Bonferroni-corrected statistical significance threshold in the sex-combined analysis (experiment-level p<1.3×10^−06^, i.e. a correction for 37,435 variants tested; **SI Appendix Table S3**).

We conducted systematic analyses of genomic context to establish whether the identified rare nonsynonymous variants are likely to be causal for the association with BMI-adjusted WHR, including conditional analyses and fine-mapping of statistically-decomposed signals (**Methods**). Fine-mapping analyses provided strong statistical evidence for the causal association of rare nonsynonymous variants of *CALCRL*, *PLIN1*, *PDE3B* and *ACVR1C* in the main analysis and of *FGF1* in the experiment-level statistical significance secondary analysis (**Table 1, Fig. 1, and SI Appendix Fig. S3**). Conversely, genomic context analyses were not consistent with causal associations for identified rare nonsynonymous variants in *ABHD15* (main analysis), *PYGM* (main analysis), *PLCB3* (experiment-level statistical significance and women-specific secondary analyses) or *FNIP1* (women-specific secondary analysis; all results in **SI Appendix Tables S3-**S4 and Note S3).

**Table 1.** Conditionally independent index variants and fine-mapping at the *CALCRL, PLIN1, PDE3B, ACVR1C* and *FGF1* loci.

| Locus | Signal | dbSNP rsID | Genomic coordinate, chromosome, position, effect allele, other allele (effect allele frequency, %) | Annotation | Beta (SE) from univariate analysis, in SD units | p-value univariate analysis | Beta (SE) from conditional analysis*, in SD units | p-value conditional analysis* | Genomic position of 99% credible set window, (width in number of base pairs) | Number of variants in the credible set | PPA for the index variant, % |
| --- | --- | --- | --- | --- | --- | --- | --- | --- | --- | --- | --- |
| *Main analysis* |  |  |  |  |  |  |  |  |  |  |  |
| *CALCRL* | 1 | rs10177093 | chr2:188213819:G:T (45.6%) | *CALCRL* intronic | -0.02 (0.002) | $2.2 \times 10^{-27}$ | -0.02 (0.002) | $7.7 \times 10^{-26}$ | 188088527-188213819 (125,293) | 60 | 17% |
| | 2‡ | rs61739909 | chr2:188245439:G:A (0.3%) | *CALCRL* p.L87P | -0.13 (0.018) | $2.0 \times 10^{-13}$ | -0.12 (0.018) | $5.9 \times 10^{-12}$ | 188245439-188270259 (24,821) | 2 | 51% |
| *PLIN1* | 1‡ | rs139271800 | chr15:90214777:G:A (0.1%) | *PLIN1* p.L90P | -0.21 (0.029) | $5.5 \times 10^{-13}$ | -0.21 (0.029) | $5.5 \times 10^{-13}$ | 90214777 (1) | 1 | >99% |
| *PDE3B* | 1‡ | rs150090666 | chr11:14865399:T:C (0.1%) | *PDE3B* p.R783X | -0.26 (0.032) | $1.4 \times 10^{-15}$ | -0.25 (0.032) | $6.2 \times 10^{-15}$ | 14865399 (1) | 1 | >99% |
| | 2 | rs2970332 | chr11:14360435:G:A (23.1%) | *RRAS2* intronic | -0.02 (0.002) | $9.9 \times 10^{-12}$ | -0.02 (0.002) | $6.3 \times 10^{-12}$ | 14258010-14689340 (431,331) | 20 | 23% |
| | 3 | rs79634051 | chr11:14561945:C:G (2.8%) | *PSMA1* intronic | -0.03 (0.006) | $6.4 \times 10^{-8}$ | -0.04 (0.006) | $2.1 \times 10^{-9}$ | 14242862-14891141 (648,280) | 15 | 78% |
| *ACVR1C* | 1 | rs55920843 | chr2:158412701:G:T (1.2%) | *ACVR1C* p.N150H | -0.08 (0.009) | $8.9 \times 10^{-19}$ | -0.09 (0.009) | $4.6 \times 10^{-20}$ | 158412701 (1) | 1 | >99% |
| | 2 | rs2444770 | chr2:158503739:C:T (14.8%) | 18kb 5' of *ACVR1C* | -0.02 (0.003) | $5.9 \times 10^{-13}$ | -0.02 (0.003) | $7.7 \times 10^{-15}$ | 158496502-158518238 (21,737) | 7 | 46% |
| | 3‡ | rs56188432 | chr2:158406865:G:A (0.2%) | *ACVR1C* p.I195T | -0.14 (0.021) | $4.9 \times 10^{-11}$ | -0.14 (0.021) | $5.4 \times 10^{-12}$ | 158406865 (1) | 1 | >99% |
| *Experiment-level p-value analysis†* |  |  |  |  |  |  |  |  |  |  |  |
| *FGF1* | 1 | rs10477191 | chr5:142077715:A:G (95.1%) | 79bp 5' of *FGF1* | -0.04 (0.005) | $5.5 \times 10^{-17}$ | -0.04 (0.005) | $7.1 \times 10^{-16}$ | 142077715 (1) | 1 | >99% |
| | 2 | rs7712968 | chr5:142086214:T:C (93.3%) | 8.6kb 5' of *FGF1* | -0.03 (0.004) | $1.3 \times 10^{-12}$ | -0.03 (0.004) | $3.5 \times 10^{-11}$ | 142082930-142101101 (18,172) | 5 | 58% |
| | 3 | rs34000 | chr5:141973501:T:C | 3'-UTR of | 0.01 | $1.2 \times 10^{-10}$ | 0.01 | $9.9 \times 10^{-8}$ | 141480886- | 192 | 30% |
|  |  |  | (59.6%) | *FGF1* | (0.002) |  | (0.002) |  | 142470008<br>(989,123) |  |  |
| | 4 | rs10065321 | chr5:141857415:C:T<br>(58.3%) | 114kb 3' of<br>*FGF1* | 0.01<br>(0.002) | $1.3 \times 10^{-9}$ | 0.01<br>(0.002) | $3.8 \times 10^{-10}$ | 141857415-141861399<br>(3,985) | 5 | 55% |
| | 5 | rs2434416 | chr5:141789701:A:C<br>(56.2%) | 85kb 5' of<br>*SPRY4* | -0.01<br>(0.002) | $8.3 \times 10^{-9}$ | -0.01<br>(0.002) | $1.5 \times 10^{-9}$ | 141769319-141824669<br>(55,351) | 34 | 22% |
| | 6<sup>‡</sup> | rs17223632 | chr5:141993631:T:C<br>(0.3%) | *FGF1*<br>p.G21E | -0.10<br>(0.018) | $8.1 \times 10^{-8}$ | -0.09<br>(0.018) | $1.6 \times 10^{-7}$ | 141589594-142413739<br>(824,146) | 62 | 98% |

Analyses are from 450,562 European ancestry individuals. Beta and standard errors are in standardized units of BMI-adjusted WHR per copy of the effect allele. Genomic coordinates according to human genome reference sequence GRCh37.
\* Adjusting for conditionally-independent index variants highlighted in the joint conditional model.
† Using a p-value threshold $p < 1.3 \times 10^{-6}$ , corresponding to a Bonferroni correction for 37,435 genetic variants studied in this analysis.
Abbreviations: SE, standard error; SD, standard deviation.
‡ Variant identified in the genome-wide scan of rare nonsynonymous variants.
Abbreviations: SE, standard error; SD, standard deviation; PPA, posterior probability of association.

**Figure 1.**
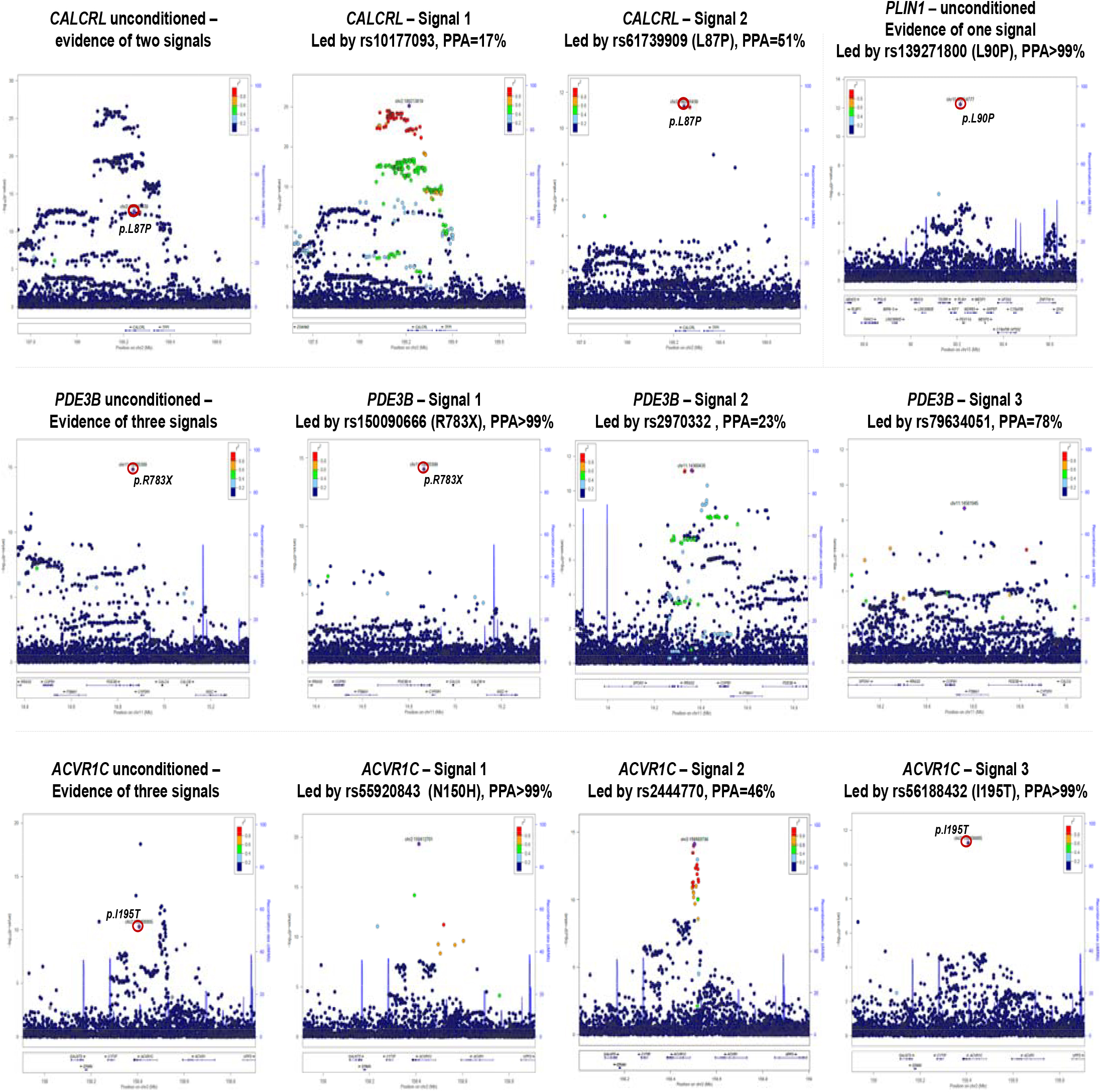
Regional association plots of the overall and statistically-decomposed signals at the *CALCRL*, *PLIN1*, *PDE3B* and *ACVR1C* genes. Plots were drawn using LocusZoom (99). Joint meta-analysis models using GCTA (79) were used at each locus to assess how many independent signals were present. Then, at each locus each signal was statistically-decomposed from others by estimating associations of all variants in the region adjusted for all other index variants at the region. Fine-mapping of each signal was performed using a Bayesian approach (80).

At *CALCRL*, there was evidence of two conditionally-independent signals (**Table 1, Fig. 1, and SI Appendix Table S5**), led by the rs10177093 common variant and by the rare p.L87P variant, respectively. Fine-mapping at the latter signal yielded a 99% credible set including only two variants, rs61739909 (*CALCRL* p.L87P, posterior probability of casual association [PPA]=51%) and rs180960888 (intronic to *CALCRL*, PPA=48.5%). Hence, p.L87P is the most likely causal variant and *CALCRL* the most likely causal gene for this signal. Previous genome-wide association studies had identified an association at this locus led by rs1569135 (6), which is in linkage disequilibrium with the lead common variant for the first signal in this larger analysis (rs10177093; R^2^=0.74). However, this association had not been linked to *CALCRL* via fine-mapping nor were nonsynonymous variants in this gene previously associated with any fat distribution phenotypes.

At *PLIN1*, there was evidence of only one signal led by the rare p.L90P variant (**Table 1 and Fig. 1**), which was the only variant in the 99% credible set (PPA>99%; **Table 1**).

At *PDE3B*, there was evidence of three signals, the strongest of which was led by the rs150090666 p.R783X nonsense variant in *PDE3B*, which was the only variant in the 99% credible set (PPA>99%; **Table 1 and Fig. 1**). As part of an analysis focused on predicted loss-of-function variants in unrelated participants UK Biobank, Emdin et al. reported an association of rs150090666 with height and, in follow-up analyses, of a combination of predicted loss-of-function *PDE3B* variants with BMI-adjusted WHR which was below the genome-wide level of significance (26). In that study, the genomic context of the association with height, but not BMI-adjusted WHR, was considered. In this study, we included a larger sample of European ancestry participants (including related individuals) and optimally accounted for relatedness and population substructure using a mixed-model, finding a genome-wide significant association for rs150090666 with BMI-adjusted WHR, with fine-mapping providing the strongest possible statistical evidence of causal association for this variant.

At *ACVR1C*, there was evidence of three distinct signals (**Table 1 and Fig. 1**). The rare p.I195T variant led one of the secondary signals at this region and was the only variant in the 99% credible set (PPA>99%; **Table 1**). In addition, the primary signal at this region was led by a low-frequency missense variant in *ACVR1C* (rs55920843, p.N150H), which also had the highest posterior probability in fine-mapping of this signal (PPA>99%; **Table 1**). Hence, fine-mapping of conditionally-independent signals at this locus converges on *ACVR1C* as causal gene for body fat distribution and p.I195T and p.N150H as causal variants for the respective association peaks.

Additional consideration of subthreshold-signals that met the experiment-level Bonferroni correction showed evidence of six conditionally-independent signals in and around the *FGF1* gene, one of which was led by the rare p.G21E missense variant (PPA=98%; **Table 1 and SI Appendix Fig. S3**).

Given previous reports of sex-specific associations with BMI-adjusted WHR (6), we estimated stratified associations for likely-causal variants identified in our analysis and, in line with previous studies (6), found larger effect-size estimates in women compared to men, with a statistically-significant difference for rs150090666 p.R783X in *PDE3B* (p_heterogeneity_=5.2×10^−06^; **SI Appendix Table S6**).

In a gene-based analysis, the burden of rare nonsynonymous alleles in *PLIN1*, the only gene with other rare nonsynonymous variants in addition to those found in the main analysis, was not associated with statistically-significant differences in BMI-adjusted WHR (**SI Appendix Table S7**).

### Functional annotation, structural modelling and associations with cardio-metabolic phenotypes

We conducted detailed *in silico* analyses that predict functional impact of identified variants (**SI Appendix Box S1 and Table S8**) and estimated their association with a variety of continuous cardio-metabolic traits and outcomes (**Fig. 2–3**), to gain insights into their likely function and phenotypic impact.

**Figure 2.**
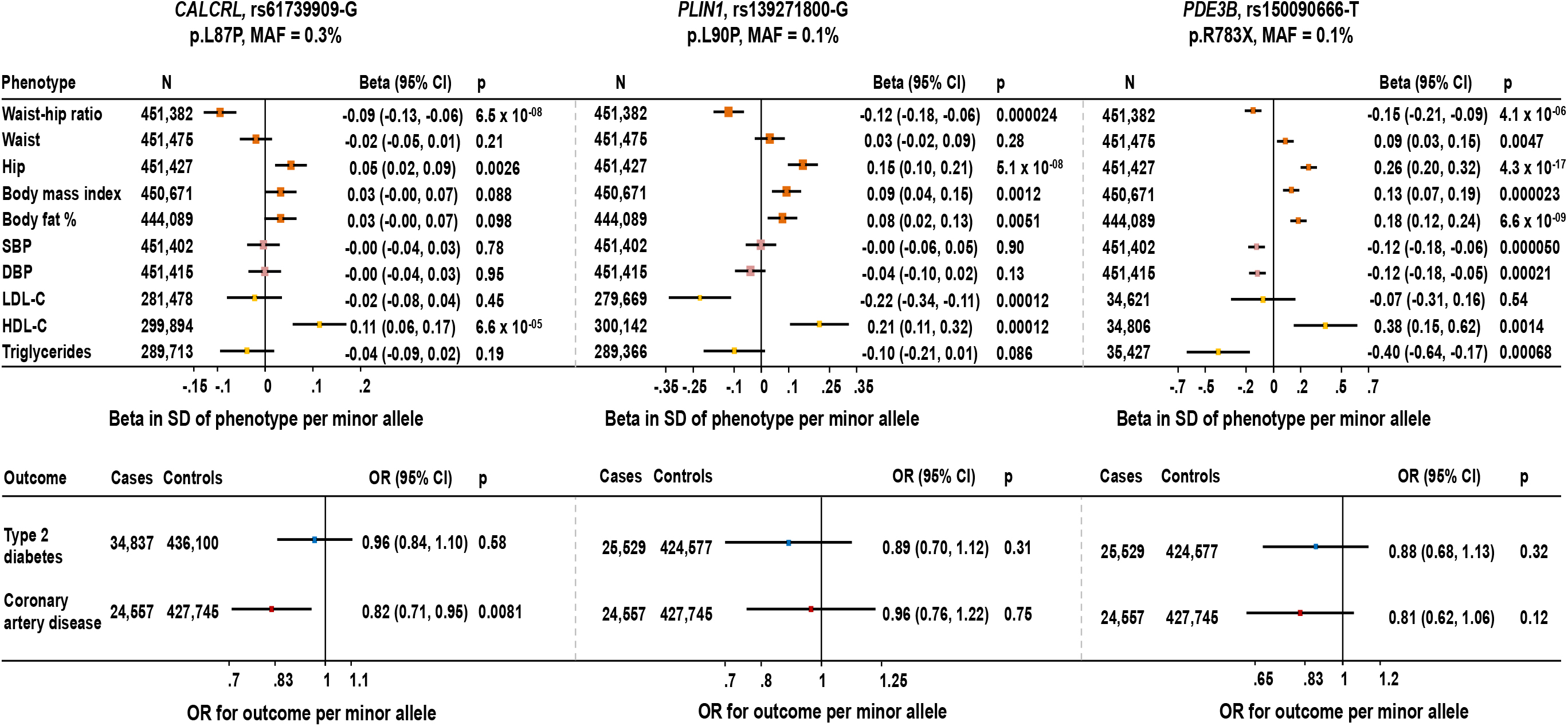
Association of rare nonsynonymous genetic variants at *CALCRL, PLIN1* and *PDE3B* with continuous metabolic traits and risk of cardio-metabolic disease outcomes. Associations are presented as beta coefficient in standardized units of continuous trait or odds ratio for disease outcome per minor allele. The minor allele is listed following the rsid above the corresponding plot. Lipid association estimates were not available for rs150090666 in the Global Lipids Genetics Consortium, so they were estimated in the Fenland, InterAct and EPIC-Norfolk studies. Abbreviations: WHR, waist to hip ratio *unadjusted* for body mass index; Waist, waist circumference; Hip, hip circumference; BMI, body mass index; BF %, body fat percentage; SBP, systolic blood pressure; DBP, diastolic blood pressure; LDL-C, low-density lipoprotein cholesterol; HDL-C, high-density lipoprotein cholesterol; SD, standard deviation; OR, odds ratio.

**Figure 3.**
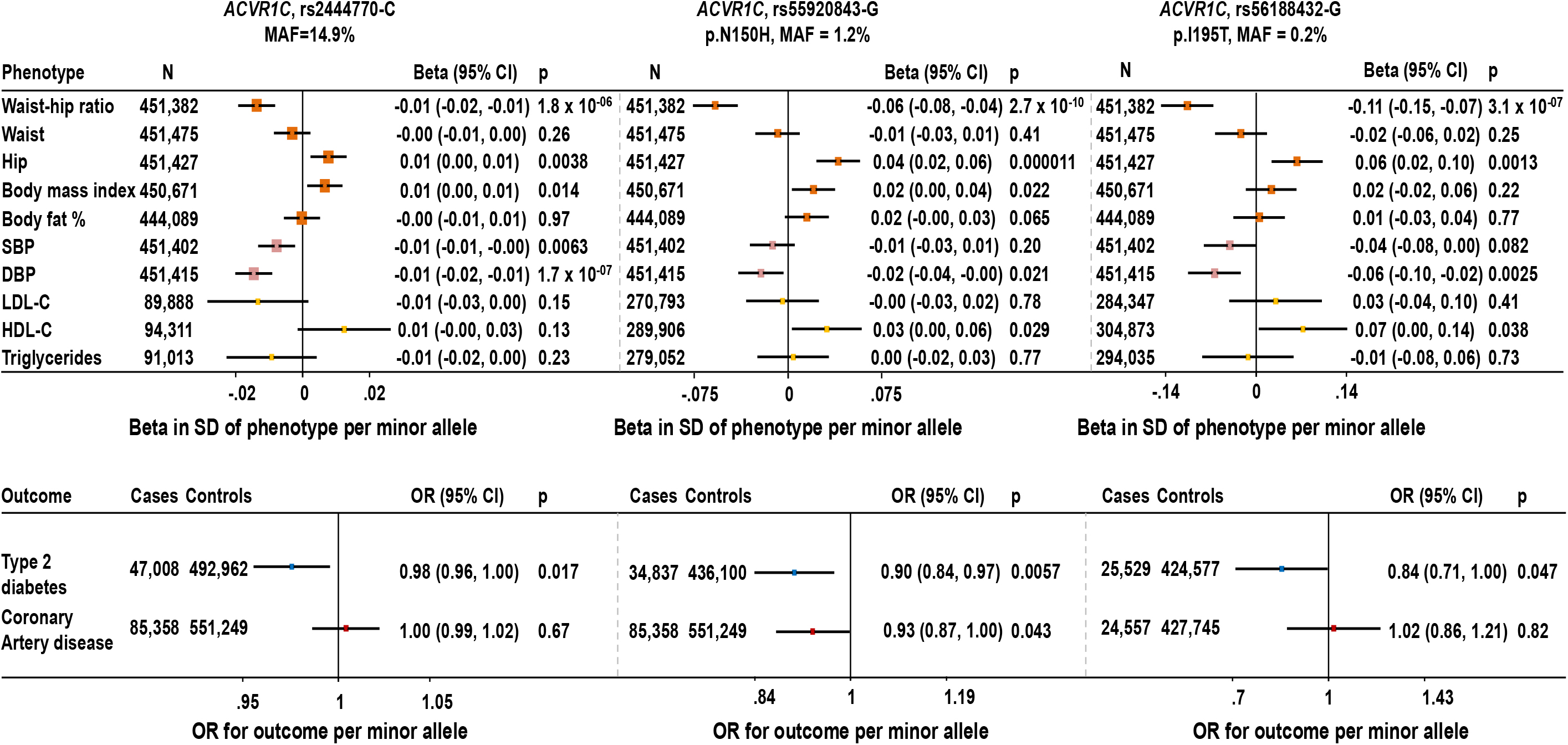
Associations with continuous metabolic traits and risk of cardio-metabolic disease outcomes of genetic variants at the *ACVR1C* gene. The three genetic variants were independently associated with waist-to-hip ratio adjusted for body mass index in conditional analyses at the *ACVR1C* gene. Associations are presented as beta coefficient in standardized units of continuous trait or odds ratio for disease outcome per minor allele. The minor allele is listed following the rsid above the corresponding plot. Abbreviations: WHR, waist to hip ratio *unadjusted* for body mass index; Waist, waist circumference; Hip, hip circumference; BMI, body mass index; BF %, body *fat* percentage; SBP, systolic blood pressure; DBP, diastolic blood pressure; LDL-C, low-density lipoprotein cholesterol; HDL-C, high-density lipoprotein cholesterol; SD, standard deviation; OR, odds ratio.

Genetic variants can affect fat distribution via increased abdominal fat (larger waist), reduced gluteofemoral fat (smaller hip) or both. The minor alleles of all four rare nonsynonymous variants in *CALCRL, PLIN1, PDE3B*, and *ACVR1C* were associated with lower waist-to-hip ratio (i.e. a more favorable fat distribution) and larger hip circumference, but were not associated with waist circumference (**Fig. 2–3**). Certain genetic variants associated with greater gluteofemoral fat show associations with protection from cardio-metabolic disease, possibly by facilitating storage in more favorable fat depots (7). At the *CALCRL, PLIN1, PDE3B*, and *ACVR1C* loci, two of the four rare nonsynonymous variants and six of nine total conditionally-independent WHR-lowering alleles were associated with protection from type 2 diabetes or coronary artery disease (p<0.05; **Fig. 2–3, SI Appendix Table S9**), suggesting that the identified variants may enhance the storage capacity of gluteofemoral adipose tissue. The four rare nonsynonymous variants were also associated with other cardio-metabolic phenotypes (**Fig. 2–3**), as described in more detail below.

The *CALCRL* p.L87P variant was associated with higher high-density lipoprotein cholesterol (HDL-C) and protection from coronary disease (**Fig. 2**). The variant occurs near a strictly conserved disulfide cross-link in this G-protein coupled receptor, but leucine 87 itself is not a conserved residue as it is replaced by proline in several species (**SI Appendix Box S1**), consistent with integrated evidence from sixteen *in silico* prediction algorithms (**SI Appendix Table S8**). This suggests that p.L87P has a mild functional impact on this G-protein coupled receptor the knock-out of which is embryonically-lethal in mice (**SI Appendix Box S1**) (27). The p.L90P variant in *PLIN1*, occurring near the conserved serine 81 which is believed to be involved in the interaction of perilipin 1 with hormone sensitive lipase (**SI Appendix Box S1**), was associated with higher overall adiposity and lower low-density lipoprotein cholesterol (LDL-C; **Fig. 2**). *In silico* prediction algorithms provide initial evidence of a likely-deleterious impact of this variant (**SI Appendix Table S8**). Loss-of-function mutations in *PLIN1* are associated with autosomal dominant forms of partial lipodystrophy with lack of gluteofemoral and leg fat, insulin resistance, dyslipidemia and type 2 diabetes (**SI Appendix Box S1**) (28). The nonsense p.R783X variant in *PDE3B* results in the premature truncation of phosphodiesterase 3B within its catalytic domain and structural modelling predicts the variant protein to be catalytically dead (**SI Appendix Box S1**). Phosphodiesterase 3B is a membrane-bound phosphodiesterase implicated in terminating intracellular lipolysis in response to insulin (29), hence an inactive enzyme would result in enhanced intracellular lipolysis at the sites where this phosphodiesterase is expressed. The variant was associated with approximately a quarter of a standard deviation lower BMI-adjusted WHR (an effect estimate six times greater than that of the strongest common variants identified in previous genome-wide association studies; **SI Appendix Fig. S4**), higher BMI and fat percentage, but lower blood pressure and lower triglycerides (**Fig. 2**). Nominal associations with protection from physician-diagnosed hypercholesterolemia and coronary artery disease have been reported for predicted loss-of-function variants in this gene (26), showing that loss-of-function of the gene may increase overall adiposity, but have protective associations with other cardio-metabolic traits (i.e. blood pressure, lipid levels, and coronary disease). In light of the statistical evidence for sex-specific association with BMI-adjusted WHR, we estimated associations of p.R783X with diabetes and coronary risk separately in men and women, but did not observe statistically-significant differences in risk for either sex (**SI Appendix Table S10**).

At *ACVR1C*, encoding a negative regulator of the peroxisome proliferator-activated receptor gamma (30), both p.N150H and p.I195T missense variants were associated with lower diastolic blood pressure and protection from type 2 diabetes (**Fig. 3**). On the basis of the three independent lead variants at this locus (**Table 1**), each standard-deviation genetically-lower BMI-adjusted WHR via *ACVR1C* was associated with a 69% lower risk of type 2 diabetes (odds ratio, 0.31; 95% confidence interval, 0.18 to 0.54; p=3.1×10^−05^). The p.I195T missense variant, which *in silico* software and structural modelling predict having a more deleterious functional impact than p.N150H (**SI Appendix Box S1 and Table S8**), also had greater phenotypic impact on WHR and diabetes risk (**Fig. 3**). At *FGF1*, we found an association for the rare WHR-lowering p.G21E allele with protection against coronary disease (**SI Appendix Fig. S5**).

Interestingly, all four genes implicated in the main analysis are abundantly expressed in subcutaneous and visceral adipose tissue in GTEx (31) and a review of functional evidence revealed links between each of the four encoded proteins and the regulation of intracellular lipolysis, the pathway responsible for the hydrolysis and release of intracellular fat from within mature cells (**SI Appendix Box S1**). Perilipin 1 and phosphodiesterase 3B are well established negative regulators of intracellular lipolysis (29, 32–34), ACVR1C has been experimentally shown to inhibit intracellular lipolysis in mouse adipocytes (30), while CALCRL is the receptor of adrenomedullin (35), which has been shown to stimulate intracellular lipolysis in human adipocytes (details in **SI Appendix Box S1**). We conducted hypothesis-free pathway-enrichment analyses using the likely-causal genes identified in this study and found evidence of enrichment for intracellular lipolysis genes (p_pathway-enrichment_=0.00093; **SI Appendix Table S11**), in addition to insulin-receptor related signaling pathways, which are established casual pathways in extreme and less severe forms of lipodystrophy (7, 36).

### Additional evidence of genetic associations at intracellular lipolysis genes

These data led us to hypothesize that variants at enzymes catalyzing the three hydrolytic reactions of intracellular lipolysis or at their direct regulators might affect fat distribution (**Fig. 4**). To test this hypothesis, we performed targeted follow-up association analyses of all genetic variation within regions 1 Mb either side of five key genes regulating each of the three hydrolytic reactions in the pathway and also estimated associations of the burden of rare nonsynonymous variants in these genes (**Methods**). While there were no associations at *G0S2* or *LIPE*, there were strong associations at *MGLL*, *ABHD5* and *PNPLA2* (p<5×10^−08^). At *MGLL* and *ABHD5* the link between genetic associations and these lipolysis genes were unclear. The association at *MGLL* was in the shadow of an association peak over 2 Mb downstream of the gene, which was greatly attenuated after conditioning for the index-variants, suggesting that this signal is unlikely to be via this gene (**SI Appendix Fig. S6 and Table S12**). At *ABHD5*, there was evidence for one association peak led by a synonymous variant in the gene (rs141365045; **SI Appendix Table S13 and Fig. S7**), which tags a low-frequency haplotype spanning the entire gene. The 99% credible set at this association signal comprises 42 variants in this haplotype that evenly share the PPA (PPA range 0.7%-3.3%), suggesting that any or a combination of these variants could be causal. The haplotype does not encompass nonsynonymous variants in the gene and the lead rs141365045 is associated with expression of the nearby *ANO10* and *SNRK-AS1* in thyroid tissue, but not *ABHD5* in GTEx (31).

**Figure 4.**
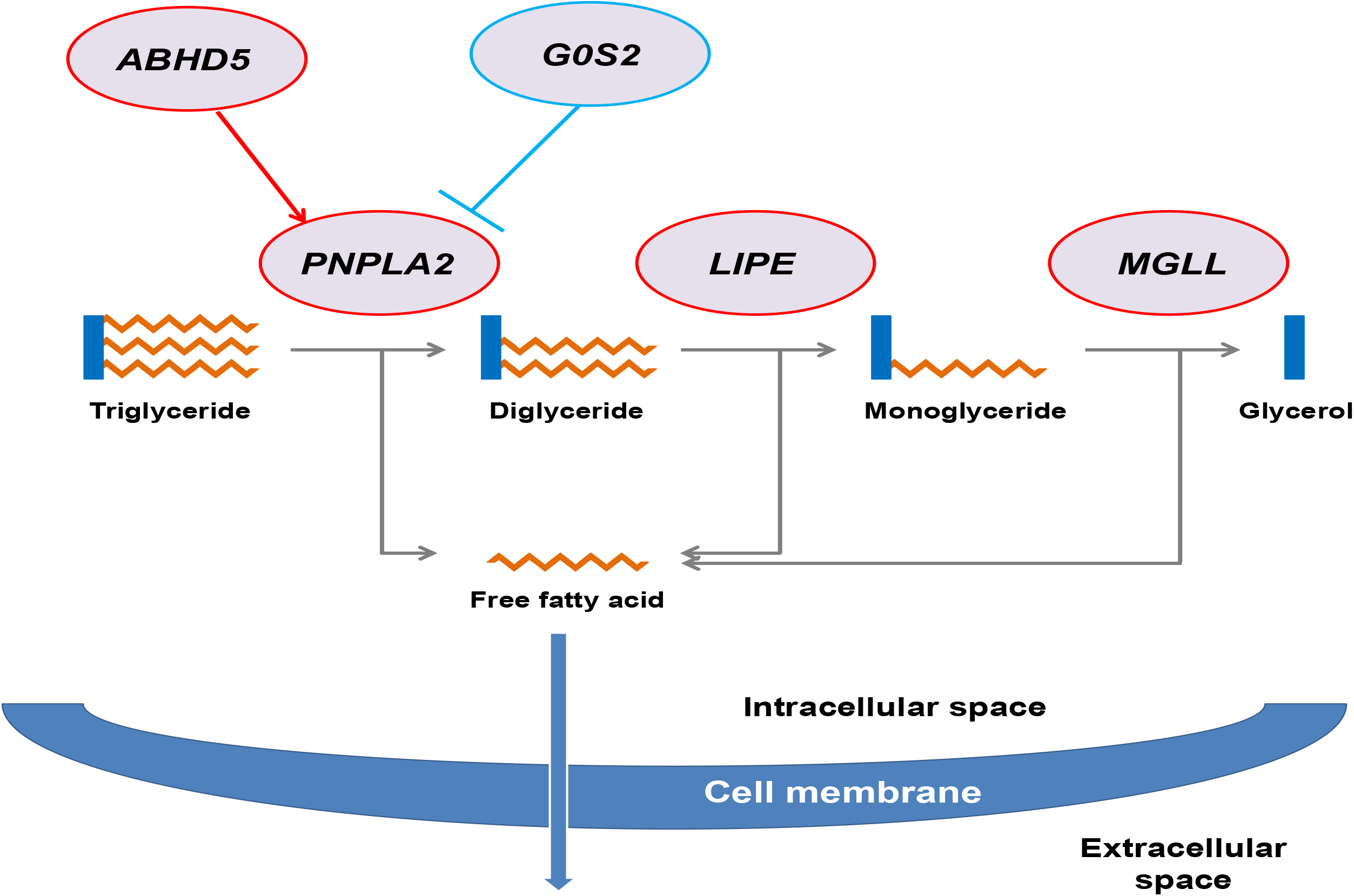
Schematic depiction of the three catalytic reactions of intracellular lipolysis and key genes in the pathway. Evidence contributing to this representation has been recently reviewed (46).

We identified an association in *PNPLA2*, encoding adipose triglyceride lipase (ATGL) which is the enzyme responsible for the initiation of intracellular lipolysis (37) (**Fig. 4**). At the locus, there was evidence of two independent signals the strongest of which was led by a missense variant occurring near a splice-site junction in the gene (rs140201358-G p.N252K; MAF=1.4%; beta in standard deviations of BMI-adjusted WHR per minor allele [252K], 0.08; standard error, 0.009; p=2.5×10^−22^; **SI Appendix Table S13 and Fig. S7**). Associations were similarly strong in men and women (p_heterogeneity_=0.10; **SI Appendix Table S6**). Fine-mapping of the main signal in the region identified rs140201358 as the only variant in the 99% credible set, supporting a likely-causal association (PPA>99%; **SI Appendix Table S13 and Fig. S7**).

Follow-up analyses of the rs140201358-G p.N252K variant showed associations with lower BMI and smaller hip circumference, but higher triglycerides and LDL-C (**Fig. 5A**). Disease outcome association analyses revealed an association with higher risk of type 2 diabetes (odds ratio per 252K allele, 1.09; 95% confidence interval, 1.02-1.17; p=0.0073) and coronary artery disease (odds ratio per 252K allele, 1.12; 95% confidence interval, 1.04-1.20; p=0.0019; **Fig. 5B**).

**Figure 5.**
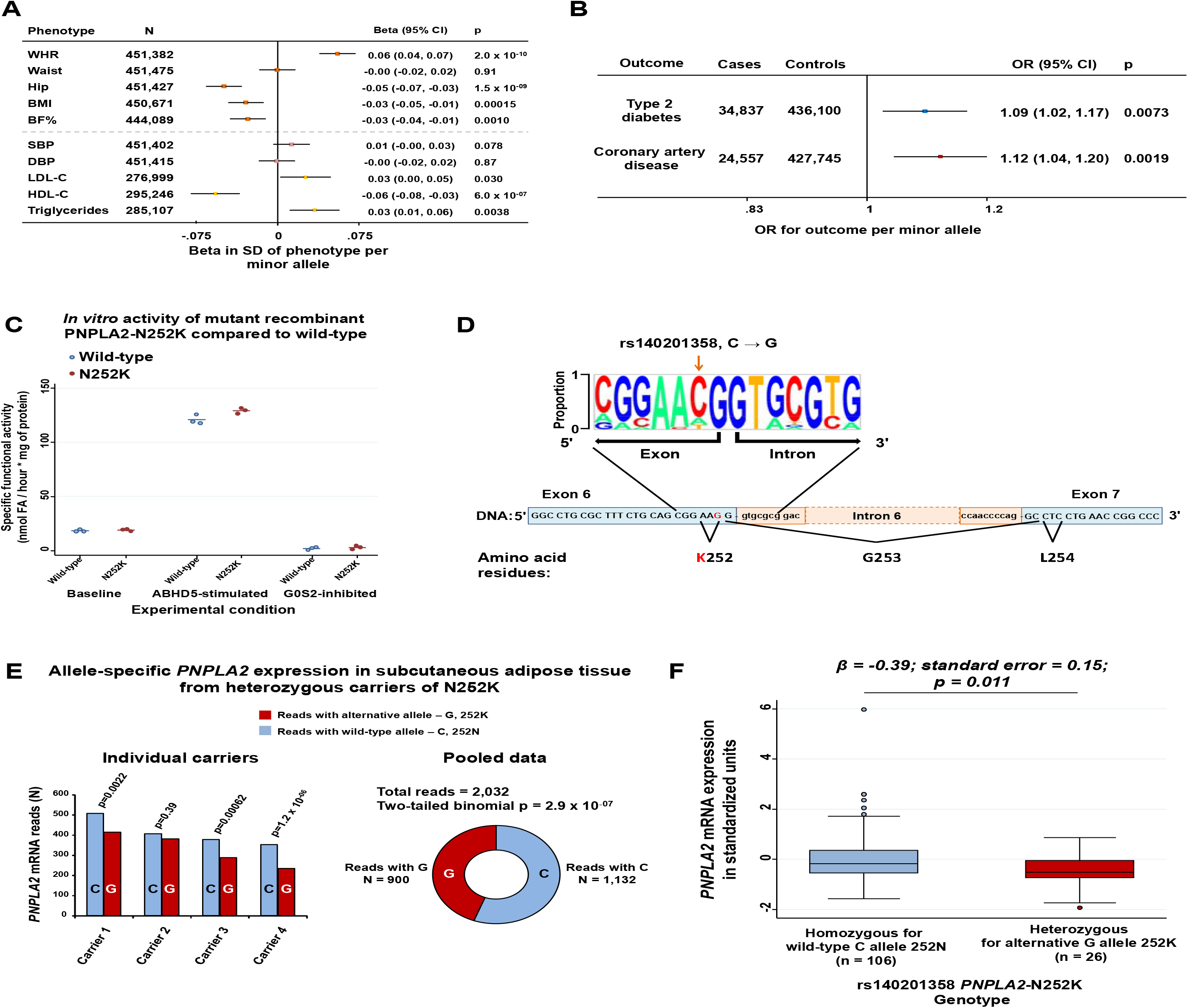
Phenotypic associations and functional consequences of the rs140201358-G variant in *PNPLA2*. ***Panel A*** reports associations with continuous traits of rs140201358-G p.N252K, while ***Panel B*** reports associations with cardio-metabolic disease outcomes. Associations are presented as beta coefficient in standardized units of continuous trait or odds ratio for disease outcome per minor allele G. The minor allele is listed following the rsid above the corresponding plot. Abbreviations: WHR, waist to hip ratio unadjusted for body mass index; Waist, waist circumference; Hip, hip circumference; BMI, body mass index; BF %, body fat percentage; SBP, systolic blood pressure; DBP, diastolic blood pressure; LDL-C, low-density lipoprotein cholesterol; HDL-C, high-density lipoprotein cholesterol; SD, standard deviation; OR, odds ratio. ***Panel C*** shows the specific enzymatic activity of PNPLA2-N252K and wild-type protein in *in vitro* expression studies. The graph reports the results of three technical experimental replicates. Full blue circles (individual replicate result) and horizontal bars (averages) are for wild type PNPLA2 while full dark red circles (individual replicate result) and horizontal bars (averages) are for p.N252K mutant PNPLA2. Abbreviations: FA, fatty acids; ABHD5, Abhydrolase Domain Containing 5 also known as Comparative Gene Identification-58 (CGI-58); G0G2, G0/G1 Switch 2. ***Panel D*** represents the location of the rs140201358 p.N252K variant at the exon 6 – intron 6 splice junction of *PNPLA2*. At the top of the panel is a representation of conservation of residues across mammalian species, with the proportion of observed nucleotides at each position represented by the size of the font. ***Panel E*** shows the results of allele-specific expression of *PNPLA2* in subcutaneous adipose tissue from four unrelated heterozygous carriers of rs140201358 p.N252K. The individual results are shown on the left, while the pooled results on the right. The reported p-values are two-tailed binomial probabilities. ***Panel F*** shows the association of rs140201358 N252K genotype with *PNPLA2* gene expression measured by quantitative transcription polymerase chain reaction in peripheral blood mononuclear cells from 106 homozygous carriers of the wild-type C allele (i.e. 252N) and 26 heterozygous carriers of the alternative G allele (i.e. 252K). The mRNA levels of *PNPLA2* were standardized on the basis of the distribution in homozygous wild-type participants. Boxes represent the median and interquartile range, whiskers represent the upper and lower adjacent values, circles represent outliers for each genotype group. Association between genotype and *PNPLA2* expression was estimated by linear regression.

We conducted a number of *in vitro* experiments to provide an initial functional characterization of the possible mechanisms linking rs140201358 with fat distribution. *In vitro* experiments showed similar basal-, ABHD5 stimulated-and GOS2-inhibited enzyme activity as well as similar localization to lipid droplets between PNPLA2-N252K and wild-type PNPLA2 (**Fig. 5C and SI Appendix Fig. S8**), in keeping with structural modelling (**SI Appendix Box S1**). However, the C>G substitution occurs at position −2 at the donor splice site of exon 6 in a partially-conserved nucleotide that is never substituted with a G in mammalian species, suggesting a possible impact on splicing (**Fig. 5D**). *In silico* software predicted this change to result in the creation of an exonic splicing silencer site, with higher probability of exon skipping (**Methods**). We hypothesized that if the variant affected the correct splicing of *PNPLA2*, this could alter allele-specific expression of *PNPLA2* in carriers. To assess this, we investigated the allele-specific expression of *PNPLA2* in subcutaneous adipose tissue from four unrelated heterozygous carriers of rs140201358-G from the Twins UK study. Across the four carriers, there were 2,032 reads of *PNPLA2* mRNA in subcutaneous adipose tissue. The number of reads carrying the alternative G allele (i.e. 252K) was 21% lower than that of reads containing the wild-type C allele (observed reads, 900; expected, 1,016; two-tailed binomial p=2.9×10^−07^), with a statistically-significant within-individual difference in three out of four carriers (p<0.05; **Fig. 5E**). To assess impact on overall expression, we conducted quantitative polymerase chain reaction (Q-PCR) analyses of *PNPLA2* expression in peripheral blood mononuclear cells from 106 homozygous carriers of the wild-type C allele and 26 heterozygous carriers of the alternative G allele from the Fenland study. Heterozygous carriers of the G allele had 0.39 standard deviations lower overall levels of *PNPLA2* mRNA compared to homozygous wild-type individuals (beta in standard deviations, −0.39; standard error, 0.15; p=0.011; **Fig. 5F**). It remains to be established if associations with expression levels do reflect a splicing defect or result from other regulatory mechanisms due to rs140201358 or correlated variants.

In gene-based analyses, the one rare nonsynonymous variant in *PNPLA2* captured by genotyping (p.S407F) or 10 rare variants in the other intracellular lipolysis genes were not associated with statistically-significant differences in BMI-adjusted WHR (**SI Appendix Table S7**).

## Discussion

By combining human genetics studies in over half a million people with *in silico* and *in vitro* functional analyses, we found evidence implicating intracellular lipolysis genes in the regulation of fat distribution and its cardio-metabolic consequences in the general population. This genetic study focused on a subset of genetic variation in order to maximize translational insights of genetic findings and is distinct from but complementary to genome-wide association studies assessing all genetic variants to clarify the overall genetic architecture of a trait. In addition, given the strict criteria for statistical significance and the systematic analysis of genomic context, all of the likely-causal alleles found in the main analysis would meet the strictest statistical significance thresholds recommended for genome-wide analyses of densely genotyped or imputed datasets, including those appropriate for the analysis of whole genome sequencing results (38). By identifying nonsynonymous alleles with high probability of being the causal variants underlying identified associations, this study provides (a) new and specific insights that go beyond the general notion of an impact of adipocyte function on body fat distribution and (b) a basis for the understanding of the molecular mechanisms behind these robust phenotypic associations. With fine-mapping, we show that five nonsynonymous variants at four of the identified genes (*PLIN1*, *PDE3B*, *ACVR1C*, and *PNPLA2*, found via targeted follow-up analysis) had >99% PPA, the highest possible statistical evidence of causal association.

Studies in rare forms of human lipodystrophy (36, 39), in experimental models (40–42) and recently also in the general population (7, 43–45) have implicated an impaired capacity to store fat in peripheral adipose compartments in cardio-metabolic disease. Our results highlight intracellular lipolysis as a novel mechanism linking impaired peripheral fat deposition to the risk of cardio-metabolic disease. Intracellular lipolysis is the biochemical process that regulates the release of fatty acid molecules from mature cells and its level of activation ultimately controls the propensity of peripheral adipocytes, and other tissues, to retain energy stores in the form of fat (46, 47). Therefore, modulating this pathway determines where and how efficiently surplus energy is stored and thus the risk of the complications of sustained positive energy balance. In addition, in a secondary analysis of this study we found evidence of a likely-causal association with lower BMI-adjusted WHR of a rare missense variant in *FGF1*, a gene that murine experiments have implicated in the remodeling of adipose tissue in response to fluctuations in nutrient availability (48). In addition to previous evidence about the role of adipogenesis and intravascular lipolysis (7), findings from this study around intracellular lipolysis and the FGF1-pathway highlight the importance of adipose tissue plasticity in response to energy availability as a critical mechanism in the determination of fat distribution and its cardio-metabolic consequences.

All four likely-causal genes identified in our hypothesis-free main-analysis of rare, nonsynonymous variants have been implicated in intracellular lipolysis by orthogonal experimental evidence. In addition, a missense variant in *PNPLA2*, encoding the initiator of intracellular triglyceride hydrolysis, was associated with unfavorable fat distribution, higher atherogenic lipid levels and higher risk of type 2 diabetes and coronary artery disease further supporting the main findings from the scan of rare variants. Rare loss-of-function mutations in *PNPLA2* cause a recessively-inherited lipid storage disease characterized by ectopic fat deposition, known as neutral lipid storage disease with myopathy, which in some of the few reported cases has been associated with dyslipidemia and diabetes (49–52). Our study is consistent with a role of intracellular lipolysis genes in the aetiology of cardiovascular and metabolic disease in the general population, in line with an earlier study of the Amish population suggesting that a deletion in hormone sensitive lipase (*LIPE*), present in ~5% of Amish people but rarely detected in other populations, results in lower intracellular lipolysis, smaller adipocytes, insulin resistance and higher diabetes risk (53).

Intracellular lipolysis is a pharmacologically modifiable pathway. The gene products of *ACVR1C* and *PNPLA2* have generated interest as potential drug targets for obesity and its complications (30, 54–57) on the basis of mouse models showing lower fat accumulation and improved glucose metabolism upon downregulation or pharmacologic inhibition of these proteins. In our human genetic studies, the peripheral adiposity-increasing alleles at these genes were associated with protection from diabetes (*ACVR1C* and *PNPLA2*) and coronary disease (*PNPLA2*). Hence, pharmacologically enhancing and not reducing peripheral fat deposition by modulating these genes could protect from cardio-metabolic disease in humans. In addition, the product of *PDE3B* is inhibited by cilostazol, a non-selective inhibitor of both phosphodiesterase 3B and 3A used in cardiovascular medicine for its anti-platelet and vasodilating properties (58, 59). The interaction between PLIN1 and ABHD5 can also be inhibited pharmacologically, resulting in enhanced PNPLA2 activity (60). The association of variation in intracellular lipolysis genes with multiple cardio-metabolic risk factors and outcomes in our study provides human genetic evidence supporting further pharmacological development for this pathway. Also, the translational implications of the association of the *FGF1* p.G21E missense variant with fat distribution and protection from coronary disease deserve further exploration in light of the mounting therapeutic interest around this pathway (61, 62).

In conclusion, our study provides human genetic evidence of a link between genes involved in the regulation of intracellular lipolysis, fat-distribution and its cardio-metabolic complications in the general population.

## Methods

### Study design and rationale

The aim of this study was to identify likely-causal nonsynonymous genetic variants and genes implicated in the regulation of body fat distribution. We performed a hypothesis-free genome-wide scan of rare (MAF<0.5% in keeping with the 1000 Genomes Project definition (23)) nonsynonymous variants coupled with systematic conditional and fine-mapping analyses at identified loci. We focused on rare variants because (a) they facilitate fine-mapping approaches for causal variant identification (25) and (b) their contribution to fat distribution is understudied (6). Since these variants are usually population-specific (23) and difficult to impute (24), their study requires large, homogeneous samples and direct genotyping. For these reasons, we focused on variants that were directly-genotyped by array genotyping in a single, large population-based cohort, the UK Biobank study (63). Analyses were focused on individuals of European ancestry. We chose to focus on nonsynonymous variation as (a) disease-associated variants are enriched for nonsynonymous variants (38), (b) if a causal variant is a nonsynonymous variant in a gene, this provides strong evidence for the causal role of the gene (64) and (c) the identification of causal nonsynonymous variants facilitates downstream functional analyses aimed at understanding the underlying mechanisms of association. An overview of the study design is in **SI Appendix Fig. S1**.

### Main and secondary analyses

In the main analysis, we studied associations of rare (MAF<0.5%) nonsynonymous variants with BMI-adjusted WHR in a sex-combined analysis of 452,302 people using the conventional threshold of genome-wide statistical significance (p<5×10^−08^). In secondary analyses, we further considered (a) subthreshold associations with BMI-adjusted WHR that met an experiment-level statistical significance threshold (p<1.3×10^−06^; corresponding to a Bonferroni correction for 37,435 tested genetic variants) or (b) genome-wide significant associations (p<5×10^−08^) in sex-specific analyses in men or women-only.

### Studies and participants

Genetic analyses were conducted in up to 452,302 European ancestry participants of UK Biobank who underwent genome-wide genotyping (**SI Appendix Table S2**). UK Biobank is a population-based cohort of 500,000 people aged between 40-69 years who were recruited in 2006-2010 from centers across the United Kingdom (63). In UK Biobank, waist and hip circumference were measured using a Seca 200cm tape measure, height was measured using a Seca 240cm measure, while weight and body fat percentage were measured using a Tanita BC418MA body composition analyzer (https://biobank.ctsu.ox.ac.uk/crystal/docs/Anthropometry.pdf). Blood pressure and resting heart rate were measured using an Omron blood pressure monitor following a standardized procedure (http://biobank.ctsu.ox.ac.uk/crystal/docs/Bloodpressure.pdf). Type 2 diabetes was defined on the basis of self-reported physician diagnosis at nurse interview or digital questionnaire, age at diagnosis > 36 years, use of oral anti-diabetic medications and electronic health records (65). Coronary artery disease was defined as either (a) myocardial infarction or coronary disease in the participant’s medical history documented by a trained nurse at the time of enrolment or (b) hospitalization or death involving acute myocardial infarction or its complications (i.e. International Statistical Classification of Diseases and Related Health Problems codes I21, I22 or I23), consistent with previous work (66, 67).

In addition to UK Biobank, genetic associations with type 2 diabetes were estimated from the EPIC-InterAct study (68) and the DIAbetes Genetics Replication And Meta-analysis (69) (DIAGRAM), with a maximum sample size of 47,008 cases and 492,962 controls. In addition to UK Biobank, genetic associations with coronary artery disease were estimated from the CARDIoGRAMplusC4D consortium (70), with a maximum sample size of 85,358 cases and 551,249 controls. Lipid traits associations were from up to 304,873 participants of the Global Lipids Genetics Consortium (71, 72). Associations with lipid traits for the p.R783X variant in *PDE3B*, which was not studied in the Global Lipids Genetics Consortium, were estimated in a meta-analysis of genetic associations in the Fenland (73), EPIC-Norfolk cohorts (74) and EPIC-InterAct subcohort (68). Descriptions of the cohorts participating in each analysis and of the sources of data are presented in **SI Appendix Table S1 and Note S1.** Ethical approvals were obtained at each study site and informed consent was obtained from all participants.

### Genome-wide association scan of rare nonsynonymous genetic variants

Similar to previous genetic studies (1, 5, 6), the BMI-adjusted WHR phenotype was constructed as the ratio of waist and hip circumferences adjusted for age, age^2^ and BMI. Residuals were calculated for men and women separately and then transformed by the inverse standard normal function. Adjustment for BMI has been suggested to possibly result in spurious associations with higher BMI-adjusted WHR of variants primarily associated with lower BMI (via collider bias) (75). However, likely-causal nonsynonymous variants at *CALCRL*, *PLIN1*, *PDE3B*, *ACVR1C, FGF1* and *PNPLA2*, were all also strongly associated with WHR not adjusted for BMI, with stronger associations than with BMI, consistent with a genuine and primary association with fat distribution (**Fig. 2–3, 5A and SI Appendix Fig. S5**). Genetic variants were genotyped in UK Biobank using the Affymetrix UK BiLEVE or the Affymetrix UK Biobank Axiom arrays (76). Genotyping underwent quality control procedures including (a) routine quality checks carried out during the process of sample retrieval, DNA extraction, and genotype calling; (b) checks and filters for genotype batch effects, plate effects, departures from Hardy-Weinberg equilibrium, sex effects, array effects, and discordance across control replicates; (c) individual and genetic variant call rate filters (76). We further excluded genetic variants with a genotype call rate below 95% and variants that were not rare or nonsynonymous. A total of 37,435 genetic variants in 12,355 genes were available for analysis. Genomic annotations were performed using the Annovar software (77). Genome-wide association analyses in 450,562 participants of European Ancestry were conducted using the BOLT-LMM software (78). BOLT-LMM fits linear mixed models that account for relatedness between individuals using a genomic relationship matrix, adjusting for relatedness and population stratification (78). Full details of these genetic analyses are in **SI Appendix Note S2**.

### Conditional analyses and fine-mapping

At each associated genomic region, we conducted systematic analyses of the genomic context of associations. Our goal was to establish whether or not the identified rare nonsynonymous variants are likely to be the causal variants for the association with BMI-adjusted WHR. At each region 1 Mb either side of the nonsynonymous genetic variants associated with BMI-adjusted WHR, we conducted both approximate and formal conditional analyses. We considered the association of all genetic variants in the regions regardless of functional annotation or allele frequency using directly-genotyped and densely-imputed data using the Haplotype Reference Consortium. First, approximate conditional analyses were conducted on summary-level estimates using GCTA (79) to identify sets of conditionally-independent index genetic variants (p<5×10^−08^ in the main or in sex-specific analyses and p<1.3×10^−06^ in analyses using experiment-level statistical significance). Individual-level genotypes for the conditionally-independent variants identified in this first step were then extracted in 350,721 unrelated European ancestry participants of UK Biobank and their independent association was confirmed in multivariable linear regression models including all variants put forward from approximate analyses. Then, at each region, we statistically decomposed the identified index signals by conditioning for the other conditionally-independent index variants. We then performed Bayesian fine-mapping (80) to estimate the posterior probability of association for each variant (PPA, where 0% indicates that the variant is not causal and 100% indicates the highest possible posterior probability that the variant is causal) and define the 99% credible set at that signal (i.e. a set of variants in a genomic window that accounts for 99% of the PPA at that association signal). To perform credible set mapping, the association results at each locus were converted to Bayes factors (BF) for each variant within the locus boundary. The posterior probability that a variant-j was causal was defined by:

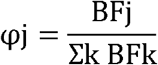

where, BFj denotes the BF for the jth variant, and the denominator is the sum of BFs for all included variants at that signal. A 99% credible set of variants was created by ranking the posterior probabilities from highest to lowest and summing them until the cumulative posterior probability exceeded 0.99 (i.e. 99%).

### Additional associations with BMI-adjusted WHR at intracellular lipolysis regulators

The findings from our rare-variant scan led us to hypothesize that variation at key enzymes of the intracellular lipolytic pathway might affect fat distribution (**Fig. 4**). To test this hypothesis, we systematically investigated variation in and around the key regulators of each of the three enzymatic reactions in intracellular lipolysis (46): *PNPLA2, ABHD5, G0S2, LIPE*, and *MGLL*. *PNPLA2* encodes adipose triglyceride lipase (PNPLA2 or ATGL), the main enzyme for triglyceride hydrolysis; *ABHD5* encodes Alpha-Beta Hydrolase Domain Containing 5 also known as Comparative Gene Identification-58 (CGI-58), the activator of PNPLA2; *G0S2* encodes G0/G1 Switch 2, the inhibitor of PNPLA2; *LIPE* encodes hormone sensitive lipase, the main enzyme for diglyceride hydrolysis; *MGLL* encodes monoglyceride lipase, the main enzyme for monoacylglyceride hydrolysis. For each of these gene regions we estimated their associations with BMI-adjusted WHR for all variants that were either directly genotyped or imputed using the Haplotype Reference Consortium in the region defined by 1 Mb either side of the gene boundaries.

### Gene-based analyses

For the four likely-causal genes identified in the main analysis (*CALCRL, PLIN1, PDE3B* and *ACVR1C*) and the five key intracellular lipolysis genes (*ABHD5*, *G0S2*, *PNPLA2*, *LIPE* and *MGLL*) we sought to estimate the association with BMI-adjusted WHR of the burden of rare nonsynonymous variants. We extracted genotypes of independent (R^2^<0.01) rare nonsynonymous variants in 350,721 unrelated European ancestry participants of UK Biobank with available BMI-adjusted WHR and estimated the burden of these variants using linear regression adjusted for age, sex and genetic principal components comparing carriers to non-carriers of these rare alleles.

### Structural modelling, functional prediction of identified nonsynonymous variants, pathway enrichment analyses

Models were built with the MODELLER software (81). Sequence alignment was achieved by HHpred, MUSCLE and Blast algorithms implemented in MPI toolkit (82). Paralogues and orthologues were extracted from Orthologous Matrix database (83), and displayed and edited in Jalview (84). Structures are displayed using MolSoft Browser-Pro software (URL: http://www.molsoft.com/icm_browser_pro.html).

We used the Annovar (77) to generate annotations that predict deleteriousness of amino acid changes. We generated the summary results of sixteen computational algorithms that predict whether or not an amino acid change is likely to be deleterious to the function of the encoded protein. For each of these algorithms, the prediction of likely functional impact contributed to an overall score of predicted deleteriousness (see **SI Appendix Table S8** for details on the algorithms and scoring criteria).

We investigated the expression of the likely-causal genes in 53 tissues from the Genotype-Tissue Expression (GTEx) consortium (31). Data were accessed from the online portal (URL: https://www.gtexportal.org/home/) on the 1^st^ of September 2017.

We performed pathway enrichment analyses using the ConsensusPathDB software (http://cpdb.molgen.mpg.de/) (85). The software integrates data from 32 public databases to identify pathways that are over-represented in a given gene list, providing a p-value for enrichment compared to what expected by chance given the number of genes in the list and the prevalence of genes of a given pathway in the list of interrogated genes (in this case the list of 12,355 genes available for analysis).

### Initial functional characterization of the PNPLA2 p.N252K variant

Given the central role of *PNPLA2* in intracellular lipolysis and the existence of established experimental protocols for studying the impact of variants of this gene on hydrolytic activity (86), we investigated the impact of the p.N252K variant. Green monkey kidney (Cos-7, ATCC and CRL-165) cells were seeded at 900,000 cells per 10 cm dish and transfected with: (a) human wild-type PNPLA2 tagged with yellow fluorescent protein, (b) human PNPLA2-N252K, and (c) LacZ as a control using Metafectene. Twenty-eight hours after transfection, cells were harvested in 300 μL HSL-buffer plus pi and disrupted by sonication. After centrifugation at 2000 g for 10 minutes at 4°C, protein concentration was determined using Bradford reagent and bovine serum albumin as a standard. Expression of human wild-type PNPLA2 and human PNPLA2-N252K was verified by Western Blotting analysis. Triglyceride hydrolase activity assay was performed as described previously (87). A total of 20 μg of Cos-7 lysates containing overexpressed human wild-type PNPLA2, human PNPLA2-N252K or LacZ as a control were incubated with 1μg purified CGI-58 (ABHD5) or 1.5 μg purified G0S2 and radiolabeled Triolein emulsified with PC/PI (0.5 mM, 20 μCi/μmol) for one hour at 37°C. Radioactivity present in the extracted fatty acids was determined using liquid scintillation counting. Activity was measured in three technical replicates, has been corrected for Cos-7 background activity (LacZ) and is presented as mean and individual results of three technical replicates.

We also investigated the intracellular localization of wild-type and mutant PNPLA2. Cos-7 cells were seeded onto coverslips in 12 well tissue culture plates with a density of 60,000 cells per well and transfected with the following constructs: (a) human wild-type PNPLA2 tagged with yellow fluorescent protein, (b) human PNPLA2-N252K, (c) human PNPLA2-S47A, which is a catalytically inactive variant, (d) human PNPLA2 with both the N252K and S47A variants. Constructs were generated by site-directed mutagenesis using the Agilent primer design software and the QuickChangeII XL Kit following manufacturer’s instructions. 400 μM oleic acid conjugated with BSA was supplemented 4 hours after transfection for 20 hours to promote lipid droplet formation. Cells were fixed with 4% formaldehyde for 15 minutes, followed by three washes in PBS, and incubated with LipidTox DeepRed 633 for 1 hour for lipid droplet staining. Cells were mounted on microscope slides with ProLong Gold Antifade Mountant and the yellow fluorescent protein-tagged PNPLA2 localization was determined using the Leica TCS SP8 confocal microscope with a 63X immersion oil objective (1.3 NA). Yellow fluorescent protein fluorescence was excited at 514 nm and emission was detected between 520 and 545 nm. LipidTox DeepRed was excited at 633 nm and detected between 640–680 nm.

The expression of yellow fluorescent protein-PNPLA2 transfected into Cos-7 cells was determined by immunoblotting as previously described (33, 86). Briefly, cells were rinsed twice with ice-cold PBS and lysed in RIPA buffer supplemented with protease inhibitors. Cell debris was spun down at 13,000 RPM for 10 minutes at 4C. Typically, 15-20 μg of the clarified lysate was resolved and transferred onto nitrocellulose membrane using the NuPAGE Bis-tris SDS PAGE/IBlot system (Invitrogen) with yellow fluorescent protein tagged-PNPLA2 being detected using an anti-GFP antibody (Roche) and GAPDH (GeneTex) serving as loading control.

### Impact of the rs140201358-G PNPLA2 variant on gene expression

Splicing consequences for the *rs140201358-G* variant were predicted using the Human Splicing Finder software (88), while the likelihood of exon skipping was predicted using the EX-SKIP software (89).

Allele-specific expression of *PNPLA2* in adipose tissue was investigated in four unrelated heterozygous carriers from 477 female participants of the TwinsUK cohort using paired whole genome sequence and RNAseq data. Phased whole-genome sequence (6X) was generated as described in the UK10K project (90). RNAseq data from subcutaneous adipose tissue was generated as described in Buil and colleagues (91). Raw RNA reads were aligned to personal genomes using the following strategy. The phased whole genome sequences from UK10K were re-aligned to the human genome build GRCh37/hg37 to create diploid personal genomes for each sample using vcf2diploid (92). RNAseq reads were processed as follows. Adapter sequences were trimmed from RNA-seq reads using TrimGalore, software that combines Cutadapt (93) and FastQC (URL: https://www.bioinformatics.babraham.ac.uk/projects/fastqc/). Trimmed sequences with less than 20 bases or which had Phred score below 1 were excluded. Poly-A tails longer than 4bp were trimmed with PRINSEQ-lite (94). Processed reads were then aligned to the corresponding personal diploid genomes using Spliced Transcripts Alignment to a Reference (STAR) (95). Pairing of RNA-seq reads was evaluated for a Mapping Quality (MAPQ) score above 30 and a maximum mismatch threshold of 5. Genomic locations of the reads of personal genomes were crosslinked to their genome locations on the reference genome with CrossMap v.0.2.3 (96). A read’s haplotype origin was assigned to either the haplotype with the least number of mismatches or assigned randomly to break ties. Uniquely mapped reads were retained and the number of reads mapping to each haplotype was quantified with ASEReadCounter (97). We tested for differential expression of the two alleles of rs140201358 (C or G) in individual carriers and in pooled data from four unrelated carriers by calculating the two-sided bionomial probability of observed reads mapping to the G allele assuming an expected probability of 0.5.

Peripheral blood mononuclear cells (PBMCs) were isolated from 1,084 participants of the population based Fenland study (7, 73) using Ficoll-Paque (VWR International Ldt) gradient centrifugation from 20 mL sodium citrate whole blood. After washing with DPBS (Sigma-Aldrich Co Ltd), cells were re-suspended in 1 mL KOSR/DMSO (Sigma-Aldrich Co Ltd) at a concentration of approximately 1×10^7^ cells/mL. Vials were frozen to −80 °C in a controlled container and then transferred to liquid nitrogen. Expression of *PNPLA2* was measured by quantitative polymerase chain reaction (Q-PCR) in 26 heterozygous carriers of the alternative G allele, constituting all carriers with available PBMCs, and a random selection of 106 homozygous carriers of the wild-type C allele. Briefly, RNA was extracted from 1-2 million PBMCs using the RNeasy Plus Micro Kit (Qiagen), following the manufacturer’s protocol. A total of 0.1 μg RNA was reverse transcribed to cDNA using M-MLV reverse transcriptase (Promega). Q-PCR was performed to determine the mRNA expression levels of *PNPLA2* and housekeeping gene hypoxanthine phosphoribosyltransferase 1 (*HPRT1*) from undiluted cDNA with TaqMan gene expression assays (ThermoFisher Scientific, Hs00386101_m1 and Hs02800695_m1, respectively), and TaqMan Universal PCR Master Mix. The mRNA expression of *PNPLA2* from each PBMC was normalized to *HPRT1* expression using a standard curve. Measures were carried out in two technical replicates. *PNPLA2* mRNA levels were standardized to a mean of 0 and a standard deviation of 1 using the distribution in wild-type homozygous carriers. The association of genotype status with *PNPLA2* mRNA levels was estimated using repeated measures general linear regression to account for duplicate measures.

### Statistical analysis

Genetic associations were estimated using linear mixed models, linear regression or logistic regression as appropriate for the outcome phenotype and study design. Results were scaled to represent the beta estimate in standardized units for continuous outcomes or the odds ratio for binary outcomes per allele. At the *ACVR1C* gene, associations of genetically-determined body fat distribution with type 2 diabetes of multiple genetic variants were estimated using an inverse variance weighted approach (98). Estimates of (1) *ACVR1C* genetic variant to BMI-adjusted WHR and (2) *ACVR1C* genetic variant to diabetes associations were used to calculate estimates of (3) genetically-higher BMI-adjusted WHR via *ACVR1C* to diabetes association. Statistical analyses were conducted using BOLT-LMM (78) and STATA v14.2 (StataCorp, College Station, Texas 77845 USA).

### Data availability

This research has been conducted using the UK Biobank resource. Access to the UK Biobank genotype and phenotype data is open to all approved health researchers (http://www.ukbiobank.ac.uk/).

#### Data download

DIAGRAM consortium (http://diagram-consortium.org/)

CARDIoGRAMplusC4D (http://www.cardiogramplusc4d.org/)

GLGC consortium (http://csg.sph.umich.edu//abecasis/public/lipids2013/; http://csg.sph.umich.edu//abecasis/public/lipids2017/)

#### Study websites

UK Biobank (http://www.ukbiobank.ac.uk/)

EPIC-InterAct (http://www.inter-act.eu/)

Twins UK (http://www.twinsuk.ac.uk/)

Fenland (http://www.mrc-epid.cam.ac.uk/research/studies/fenland/)

EPIC-Norfolk (http://www.srl.cam.ac.uk/epic/)

#### Online data or software

Human Splicing Finder (http://www.umd.be/HSF3/)

EX-SKIP (http://ex-skip.img.cas.cz/)

FastQC (https://www.bioinformatics.babraham.ac.uk/projects/fastqc/)

MolSoft Browser-Pro (http://www.molsoft.com/icm_browser_pro.html)

ConsensusPathDB (http://cpdb.molgen.mpg.de/)

Genotype-Tissue Expression (GTEx) consortium (https://www.gtexportal.org/home/)

## Acknowledgement

This research has been conducted using the UK Biobank resource. Access to the UK Biobank genotype and phenotype data is open to all approved health researchers (http://www.ukbiobank.ac.uk/). This study was funded by the United Kingdom’s Medical Research Council through grants MC_UU_12015/1, MC_PC_13046, MC_PC_13048 and MR/L00002/1. This work was supported by the MRC Metabolic Diseases Unit (MC_UU_12012/5) and the Cambridge NIHR Biomedical Research Centre and EU/EFPIA Innovative Medicines Initiative Joint Undertaking (EMIF grant: 115372). Data from the EPIC-InterAct study contributed to this study. EPIC-InterAct Study funding: funding for the InterAct project was provided by the EU FP6 programme (grant number LSHM_CT_2006_037197). D.B.S. and S.O’R. are supported by the Wellcome Trust (WT107064 and WT 095515 respectively) the MRC Metabolic Disease Unit, the National Institute for Health Research (NIHR) Cambridge Biomedical Research Centre and the NIHR Rare Disease Translational Research Collaboration. K.S.S. is supported by MRC Project Grant L01999X/1. The TwinsUK study was funded by the Wellcome Trust and European Community’s Seventh Framework Programme (FP7/2007-2013). The TwinsUK study also receives support from the National Institute for Health Research (NIHR)-funded BioResource, Clinical Research Facility and Biomedical Research Centre based at Guy’s and St Thomas’ NHS Foundation Trust in partnership with King’s College London. Some computation was enabled through access granted to K.S.S. to the MRC eMedLab Medical Bioinformatics infrastructure, supported by the Medical Research Council (grant number MR/L016311/1). M. I. McC. is a Wellcome Senior Investigator supported by Wellcome grants 098381, 090532, 106130, 203141. M. I. McC. was supported by the National Institute for Health Research (NIHR) Oxford Biomedical Research Centre (BRC) and the views expressed in this article are those of the author(s) and not necessarily those of the NHS, the NIHR, or the Department of Health. Dr. R. A. S. is an employee and shareholder of GlaxoSmithKline Plc. (GSK). The authors gratefully acknowledge the help of the MRC Epidemiology Unit Support Teams, including Field, Laboratory and Data Management Teams.

